# TAZ/TEAD complex regulates TGF-β1-mediated fibrosis in iPSC-derived renal organoids

**DOI:** 10.1101/2021.04.15.440011

**Authors:** Xiaoping Yang, Marco Delsante, Parnaz Daneshpajouhnejad, Paride Fenaroli, Kira Perzel Mandell, Xiaoxin Wang, Shogo Takahashi, Marc K. Halushka, Jeffrey B. Kopp, Moshe Levi, Avi Z. Rosenberg

**Author notes:** **<u>Corresponding author</u>** Avi Rosenberg, MD, PhD, Department of Pathology, Johns Hopkins Medicine, Ross Building Room 632D, 720 Rutland Avenue, Baltimore, Maryland, 21205, USA.

## Abstract

Chronic kidney disease (CKD) progresses by replacement of functional tissue compartments with fibrosis, representing a maladaptive repair process. Shifting kidney repair towards a physiologically-intact architecture, rather than fibrosis, is key to blocking CKD progression. In this study, we developed a fibrosis model that uses human induced pluripotent stem cell (iPSC)-based three-dimensional renal organoids, in which exogenous TGF-β1 induces production of extracellular matrix. In these organoids, TGF- β1 increased transcription factor tafazzin (TAZ) expression. Further, in human kidney biopsies, nuclear TAZ expression was markedly increased in mild and moderate fibrosis. In cultured renal tubular cells expressing a fibrogenic program, TAZ formed a trimeric complex with phosphorylated mothers against decapentaplegic homolog 3 (p-SMAD3) and TEA domain protein (TEAD)-4. Overexpression of TEAD4 protein suppressed collagen-1α1 (*COL1A1*) promoter activity, and expression of TAZ attenuated this inhibition. INT-767, a dual bile acid receptor agonist binding farnesoid X receptor (FXR) and the Takeda G protein-coupled receptor 5 (TGR5), decreased the TGF-β1-induced increase in p-SMAD3 and TAZ, and preserved renal organoid architecture. These data demonstrate, in an iPSC-derived renal organoid fibrosis model, that INT767 prevents fibrosis programs early in the course of tubular injury through modulation of the TEAD4/TAZ pathway.

## Introduction

Chronic kidney disease (CKD) is the ninth leading cause of death in the United States and affects 37 million people, representing over 10% of the population (1). CKD can progress to end-stage kidney disease (ESKD), with substantial morbidity and mortality (2). Progressive glomerular and tubular injury are accompanied by interstitial fibrosis, a process of maladaptive repair following kidney injury. Progressive fibrosis compromises vascular function and distorts and ultimately destroys nephron architecture (2). Shifting kidney repair away from fibrosing mechanisms and toward stabilization and restoration of normal cellular and functional phenotypes are critical steps toward halting CKD progression.

To date, much of the research into the mechanisms of renal fibrosis has been performed in rodent models (3) and in two-dimensional cell culture systems (4). Although experimental animal models have provided valuable information, their genome, anatomy and physiology differ from those of humans. Furthermore, animal experiments are time consuming, the results may be strain-dependent, and for some, are ethically problematic. In many cases, the conclusions are difficult to translate to human diseases. An alternative approach is a two-dimensional cell culture system but this generally lacks relevant anatomic and physiological cues and does not replicate organ function. Nevertheless, these approaches can involve the use of human cells and the expression of human genes and proteins and reduce dependence on animal research. We sought to improve on current approaches and to do so in way that would advance the goals of personalized medicine. Our approach was to establish a relatively rapid *in vitro* model using pluripotent cell lines, that are differentiated to form three-dimensional organotypic structures, in order to study renal fibrosis mechanisms, identify patient-specific therapies, and screen for novel therapeutics

Transforming growth factor (TGF)-β is a potent driver of fibrosis (5). In the canonical pathway, TGF-β1 binds cell-surface receptor tyrosine kinases, phosphorylating mothers against decapentaplegic homolog 2 and 3 (SMAD2/SMAD3). Together with phosphorylated SMAD4 (p-SMAD4), these proteins translocate to the nucleus and activate pro-fibrotic genes. In addition to the TGF-β1-SMAD signaling axis, signaling pathways such as WNT and Hippo also contribute to maladaptive tissue repair and fibrosis (6, 7) with extensive crosstalk between TGF-β1 and Hippo signaling (7–9). Many studies of fibrogenesis and therapeutics have addressed drivers rather than mediators of TGF-β1 signaling; the latter include downstream transcription factors and their interactions (10).

Although the TGF-β pathway is a central contributor to many fibrosing disorders (5), several anti-fibrotic drugs targeting this pathway have failed in clinical trials, although the reasons for this remain unclear (11, 12). Other fibrosis pathways containing potentially novel druggable targets are being investigated, including bile acid signaling pathways. Recently, obeticholic acid, a farnesoid X receptor (FXR) agonist, has shown antifibrotic efficacy in a clinical trial of chronic liver disease (13). INT-767 is a dual bile acid receptor agonist, activating both FXR and Takeda G protein-coupled receptor 5 (TGR5). INT-767 prevents progression of diabetic and obesity-related nephropathy in mice, and also partially reverses murine age-related kidney disease (14, 15). In these models, INT-767 decreases proteinuria, podocyte injury, TGF-β1 expression and fibronectin accumulation, thus preserving renal architecture and functioning.

In the present study, we used iPSC-derived kidney organoids and renal tubular epithelial cell lines to gain further insight into the mechanism by which INT-767 blocks fibrogenesis. We demonstrate that p-SMAD3, tafazzin (TAZ) and TEA domain protein 4 (TEAD4) participate in crosstalk in the regulation of pro-fibrotic gene expression, a process that was inhibited by INT-767. The relevance of renal epithelial TAZ expression was confirmed in human kidney biopsies, wherein renal tubular epithelial TAZ expression was increased in cases of mild and moderate fibrosis. Notably, we found that INT-767 effectively inhibits TGF-β1-driven organoid fibrosis and preserves mature nephron architecture. Thus, we demonstrate the utility of iPSC-derived kidney organoids for fibrosis research to dissect relevant pathways and to screen for effective small molecular inhibitors.

## Results

### Kidney organoids were generated from human iPSCs

Human iPSCs (generous gift from Dr. Liming Gou, Johns Hopkins University, Baltimore MD, (16)) were subjected to a modified Takasato protocol to generate renal organoids (**Figure 1, Supplemental Table 1**). The time course is shown in **Figure 1** (top panel). CHIR99021, an inhibitor of glycogen synthase kinase 3 (GSK3), was added to differentiation medium (APEL2, Stem Cell Technologies, Cambridge, MA) to induce WNT signaling, as previously reported (17). After aggregating in culture with fibroblast growth factor 9 (FGF9), differentiated mesoderm cells were allowed to self-organize to form kidney organoids. Further experiments were performed at culture day 25.

**Table 1:** TAZ staining intensity in kidney biopsies with fibrosis. Table 1 summarizes Taz staining intensity in 44 renal biopsies. Cases of mild/moderate tubulointerstitial scarring have on average the most Taz intense staining, with a decrease in staining in severe scarring.

|  | TAZ +++ | Total | TAZ+++/Total |
| --- | --- | --- | --- |
| No/ Minimal | 1 | 7 | 14% |
| Mild/Moderate | 10 | 26 | 38% |
| Severe | 2 | 11 | 18% |

**Figure1.**
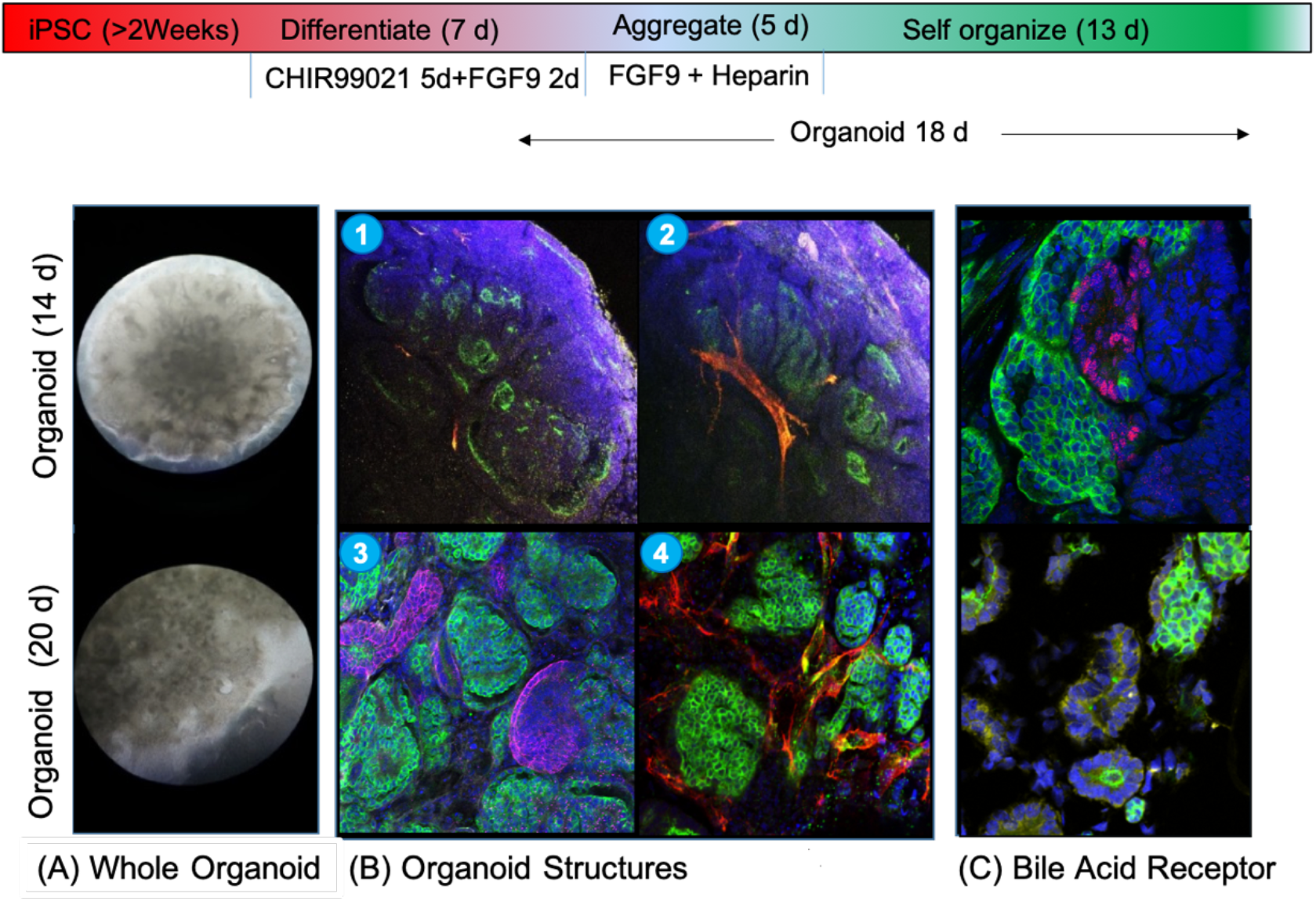
Kidney organoid generated from human iPSC. The upper panel illustrates the timeline for using iPSC to develop kidney organoids. Recovered frozen iPSCs were maintained in complete E8 medium for at least two weeks and through 3-4 passages. Viable iPSC were plated for differentiation over 7 d, during which time they received the GSK-3 inhibitor CHIR99021 for 5 d and FGF9 plus heparin for 2 d (days 6 and 7). Cells aggregated for 5 d and self-organized for 13 d, for a total 25 d required for organoid generation. The lower panel shows three-dimensional kidney organoids. (A). Whole kidney organoid, brightfield image at 14 days (*upper panel*) and at 20 days (*lower panel*). (B) Renal parenchymal compartments in whole organoids at 20 days. Podocytes express podocalyxin, PODXL (green); vascular endothelium expresses CD31 (PECAM1, platelet endothelial cell adhesion molecule), red. (10x, 2 photon microscopy) (*upper subpanel 1 and 2*). Renal parenchymal compartments in whole organoids include podocyte (expressing PODXL, green, *subpanel 3 and 4*), tubules (expressing E-cadherin purple, *subpanel 3*), and endothelium (expressing CD31, red, *subpanel 4*). Image 20x, obtained using 2-photon microscopy (*lower panel 3 and 4*). (C) Bile acid receptor expression in renal organoids. FXR-red (*upper*), TGR5 –yellow *(lower*) (40X).

Mature kidney organoids had a diameter of 5-7 mm and manifested a three-dimensional architecture (**Figure 1A**). Multiple cell lineages were demonstrated. These included podocytes (expressing podocalyxin and nephrin) in primitive glomerular structures, epithelium-lined tubules expressing proximal tubule markers (E-cadherin and *Lotus tetragonolobus* lectin) and an endothelium-lined vascular network penetrating glomerular structures (**Figure 1B**). The organoids also expressed the bile acid receptor markers of FXR and TGR5, exclusively in the tubular epithelial compartment (**Figure 1C**).

### Dose-dependent TGF-β1-induced kidney organoid fibrogenesis

To establish a kidney organoid fibrosis model, we selected TGF-β1, a potent pro-fibrotic factor, to induce fibrogenesis within the ‘mature’ organoid. We found that 5 ng/mL of TGF-β1 induced extracellular matrix (ECM) elaboration within 48-72 hours in organoids, as evidenced by increased ‘interstitial’ collagen 1-α1 (COL1A1) (**Figure 2A**) and thickened structures resembling tubular basement membranes (expressing laminin α1) (**Figure 2B**). The extent of extracellular matrix increased in a TGF-β1 dose-dependent manner and at the highest concentration (20 ng/mL), there was complete disruption of renal organoid architecture (**Figure 2A**).

**Figure 2.**
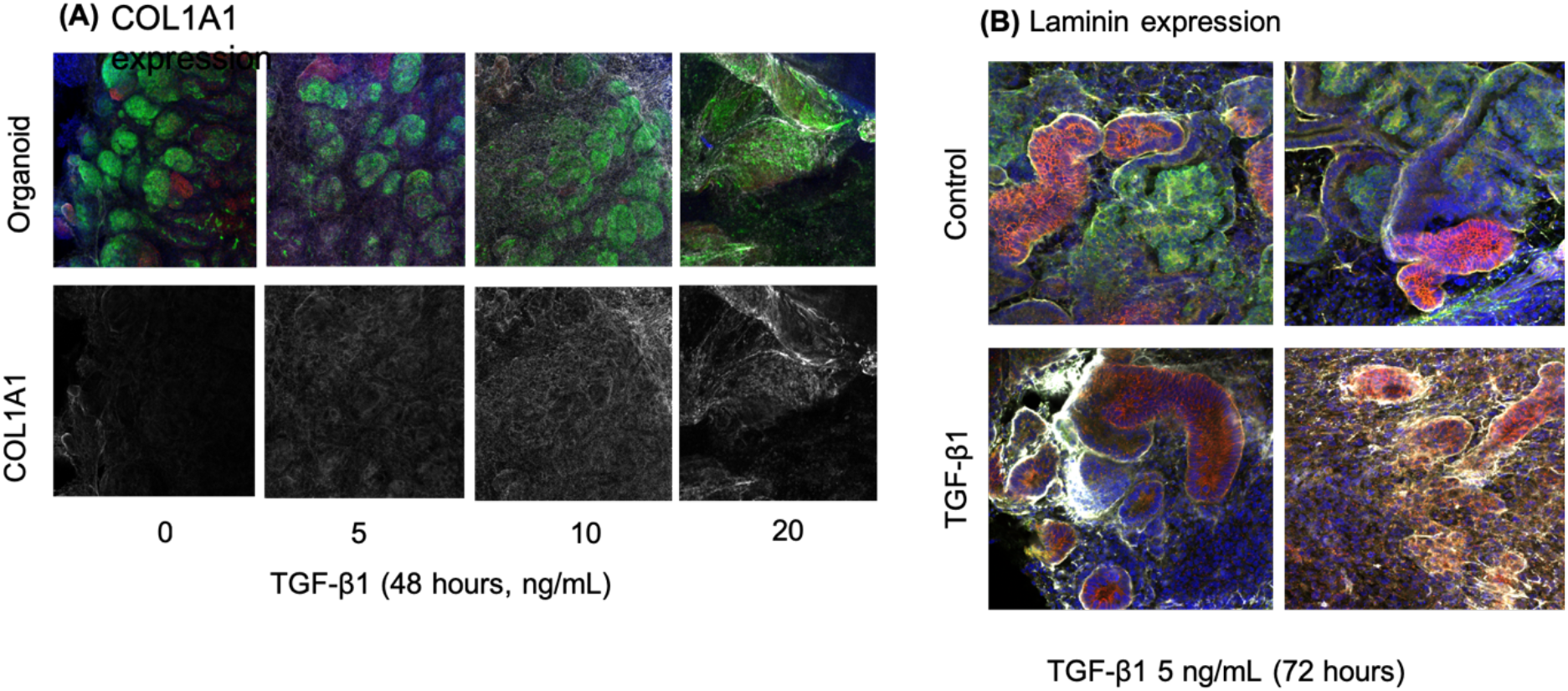
TGF-β1 induced kidney organoid fibrosis. (A) TGF-β1 induced collagen 1α1 expression in a dose-dependent manner. TGF-β1 at a high concentration (20 ng/mL) progressively disrupted nephron architecture. Glomeruloid structures expressed podocalyxin (green) and tubules expressed E-cadherin protein (red) and collagen I (α1) chain protein (COL1A1) (gray). (B) TGF-β1 induced thick basement membranes and increased interstitial laminin, in parallel with decreased expressed of proximal nephron markers (nephrin) Compared with control organoids (upper panels). TGFβ1-treated organoids (lower panels) showed decreased podocyte marker expression (nephrin, green), preserved, but dilated tubules (E-cadherin, red) and increased laminin α1 expression (gray).

### Dual bile acid receptors FXR/TGR5 agonist INT-767 decreased TGF- β1 induced ECM and preserved organoid architecture

Using this kidney organoid fibrosis model, we tested several potential anti-fibrotic compounds, including the specific inhibitor of pSMAD3 (SIS3); an inhibitor of rho-associated protein kinase (ROCK), Y27632 (data not shown); and a dual bile acid receptor agonist, INT-767. INT-767 (20 ng/mL) reduced 70% of TGF-β1-induced ECM production (COL1A1, encoding collagen 1 alpha 1 chain) and preserved “mature” kidney organoid architecture (**Figure 3A and B**). These results demonstrate that iPSC derived organoids can be used to screen for antifibrotic agents.

**Figure 3.**
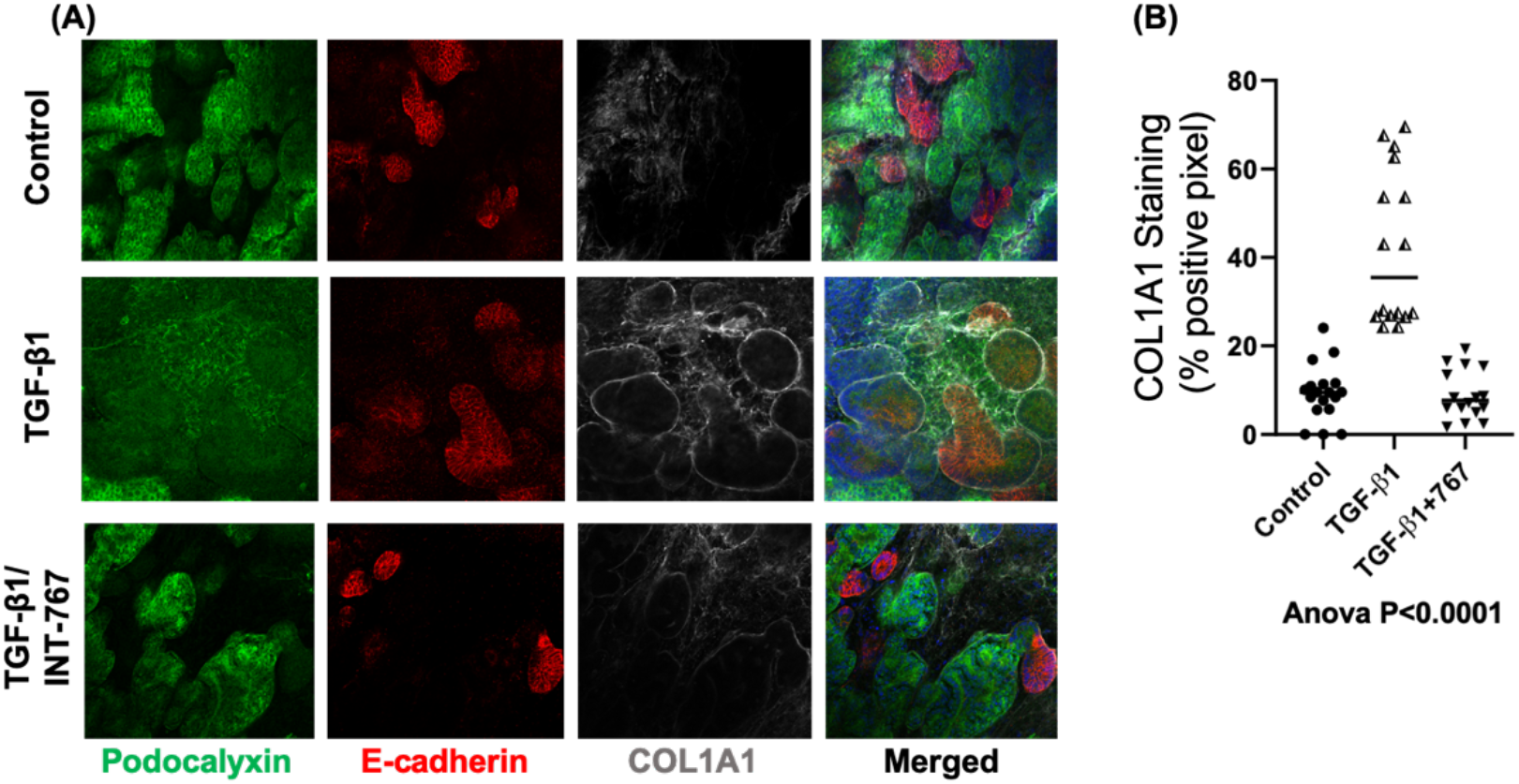
Bile acid receptor agonism ameliorates TGF-β1-induced fibrogenesis in renal organoids. (A) Dual bile acid receptor/TGR5 agonist INT-767 decreased TGF-β1-induced collagen α1 chain expression, prevented nephrin loss, and preserved normal glomerular structure. Control, TGF β1 exposure (5 ng/ mL); TGF-β1 plus INT767 (5 ng/mL/20µM respectively). (B) Quantitation of collagen1α1 (percent total area). COL1A1 dye % in quantitated area was quantified including 5 organoids for each group, and each organoid has 4-5 image sections were analyzed. ANOVA test among group p<0.0001. Two pairwise comparisons were made, Control vs TGF-β1 treated and TGF-β1 treated vs TG-β1/INT767 treated, both p <0.05.

### INT-767 increased FXR expression and suppressed p-SMAD3 expression in organoid renal tubules

It has been reported that INT-767 protects against mouse kidney injury associated with diabetes, obesity and aging (13, 15). However, the molecular mechanism by which the drug blocks fibrosis remains to be fully defined. To explore the role of INT-767 in suppressing a pro-fibrotic transcription program, we assessed expression of FXR, the INT-767 target receptor. When kidney organoids were exposed to TGF-β1 for 48 h, FXR protein expression in tubular epithelial nuclei was decreased and this TGF-β1-mediated effect was blocked by INT-767 (**Figure 4A**). However, expression of TGR5 protein (also known as G-protein coupled bile acid receptor 1), the other INT-767 target, remained unchanged (Data no shown). Using phosphorylation of SMAD3 as a marker of TGF-β1 activity, we found induction with TGF-β1 treatment that was abrogated by INT-767 co-treatment (**Figure 4B**). By immunofluorescence staining, we confirmed that the TGF-β1-induced p-SMAD3 expression occurred within renal tubular epithelium, with nuclear localization, and this expression was blocked by pre-treatment with INT-767 (**Figure 4B left)**.

**Figure 4.**
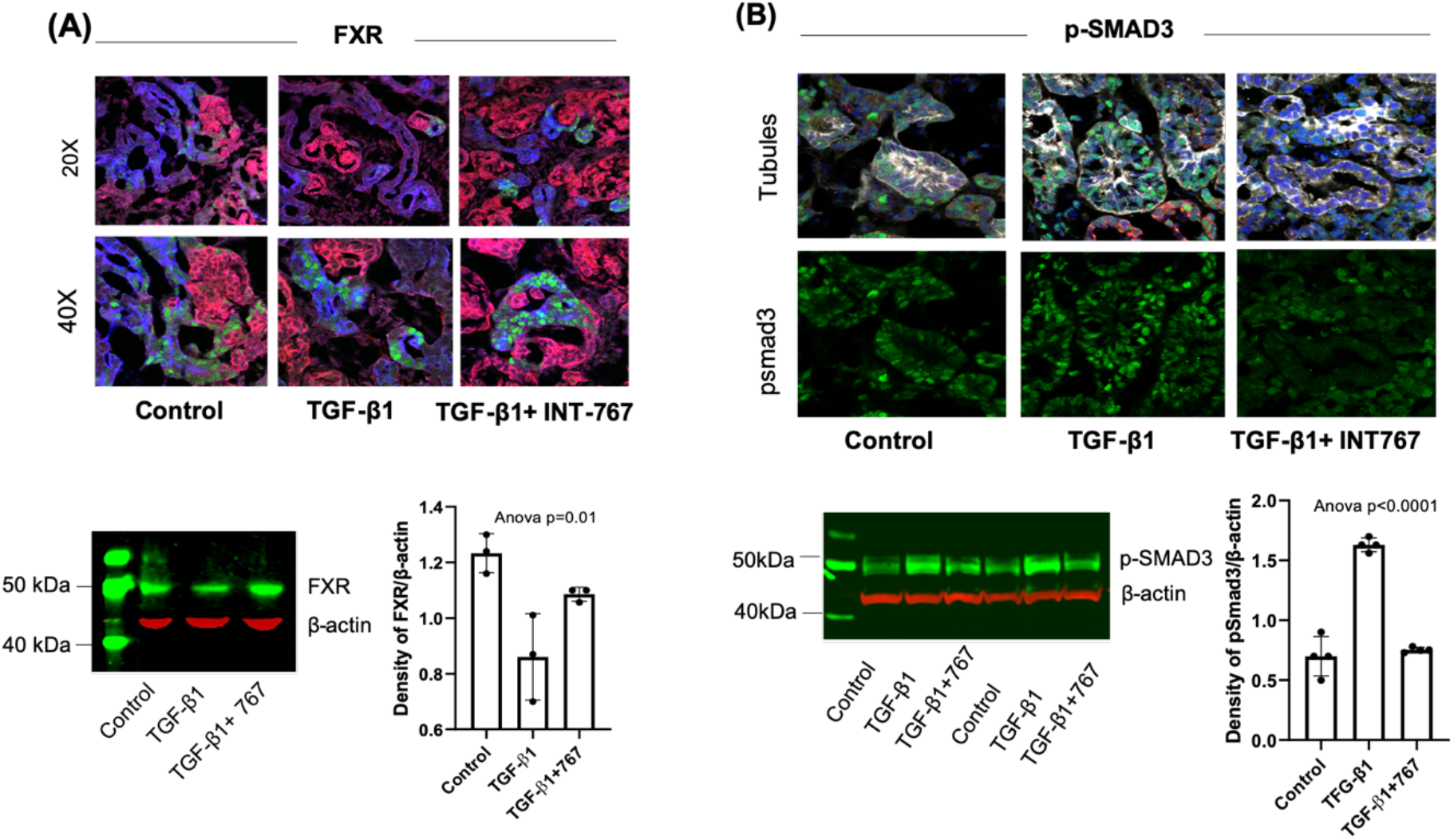
Decreased FXR and increased p-SMAD3 expression in TGF-β1 exposure. (A) TGF-β1 increased FXR protein expression in organoids. After TGF-β1 exposure for 48 h, renal organoid showed decrease of FXR staining compared to control. Addition of INT-767 increased FXR expression; the FXR was located in renal tubular nuclei (blue, Lotus tetragonolobus lectin tubule staining; red, nephrin (glomeruli); green FXR. Upper panel 20X magnification, lower panel 40X, two photons. Organoid immunoblotting showed decrease of FXR in TGF-β1-exposed samples, while there was increased FXR expression in INT-767 exposed samples (low panels, n=3 right). (B) TGF-β1 induced increase in p-SMAD3. TGF-β1 exposure for 48 h increased accumulation of p-SMAD3 in tubular nuclei, while addition of INT-767 decreased the accumulation (left panel); gray, ECAD; red, nephrin; blue, DAPI; green, p-SMAD3. 40X magnification. Organoid immunoblotting confirmed increased p-SMAD3 in TGF-β1 treatment and decreased p-SMAD3 with INT-767 treatment (low panels). Density ratio of p-SMAD3 per β-actin were pooled from three independent experiments (n=4) and shown as mean ± SD. Groups were compared by ANOVA p<0.0001. Two pairwise comparisons, Control vs TGF-β1 treated and TGF-β1 treated vs TG-β1+INT767 treated, both p <0.0001.

We extended these findings to HK2 immortalized human proximal renal tubular cells. Following TGF-β1 exposure, p-SMAD3 was progressively increased over a 72-hour time course, and pre-treatment with INT-767 attenuated this increase (**Supplemental Figure 1)**. These data support the conclusion INT-767 antagonizes TGF-β1 signaling in FXR-expressing renal tubular epithelial cells.

### INT-767 attenuated TGF- β1 induced expression of the transcriptional co-activator TAZ and profibrotic transcriptional program

INT-767 is an agonist for the bile acid receptor TGR5, a G-protein coupled receptor (GPCR), as well as for FXR. GPCR may act via Hippo signaling to promote activation of transcriptional co-activator YAP1 or its paralog TAZ (18), as well as nuclear binding proteins TEADs (19). The YAP/TAZ-TEAD axis may enhance or repress fibrotic gene transcription, depending on cellular context. Furthermore, there is cross talk between TGF*- β*1 and Hippo signaling in fibrogenesis (7).

We hypothesized that INT-767 may affect the GPCR-YAP/TAZ-TEAD axis. Therefore, we evaluated YAP and TAZ expression in relation to exposure to TGF-β1 and to TGF-β1 plus INT-767. We found that TGF- β1 increased expression of TAZ and fibrosis-related mRNAs. Interestingly, the dual bile acid receptor agonist INT-767 prevented these increases (**Figure 5B**). Immunoblotting showed that TAZ but not YAP1 was increased in TGF-β1-treated organoids. Addition of TGF-β1 increased TAZ protein by 60%, while that increase was diminished by 20% with the combination of INT-767 and TGF-β1 (p<0.0001) (**Figure 5A**).

**Figure 5.**
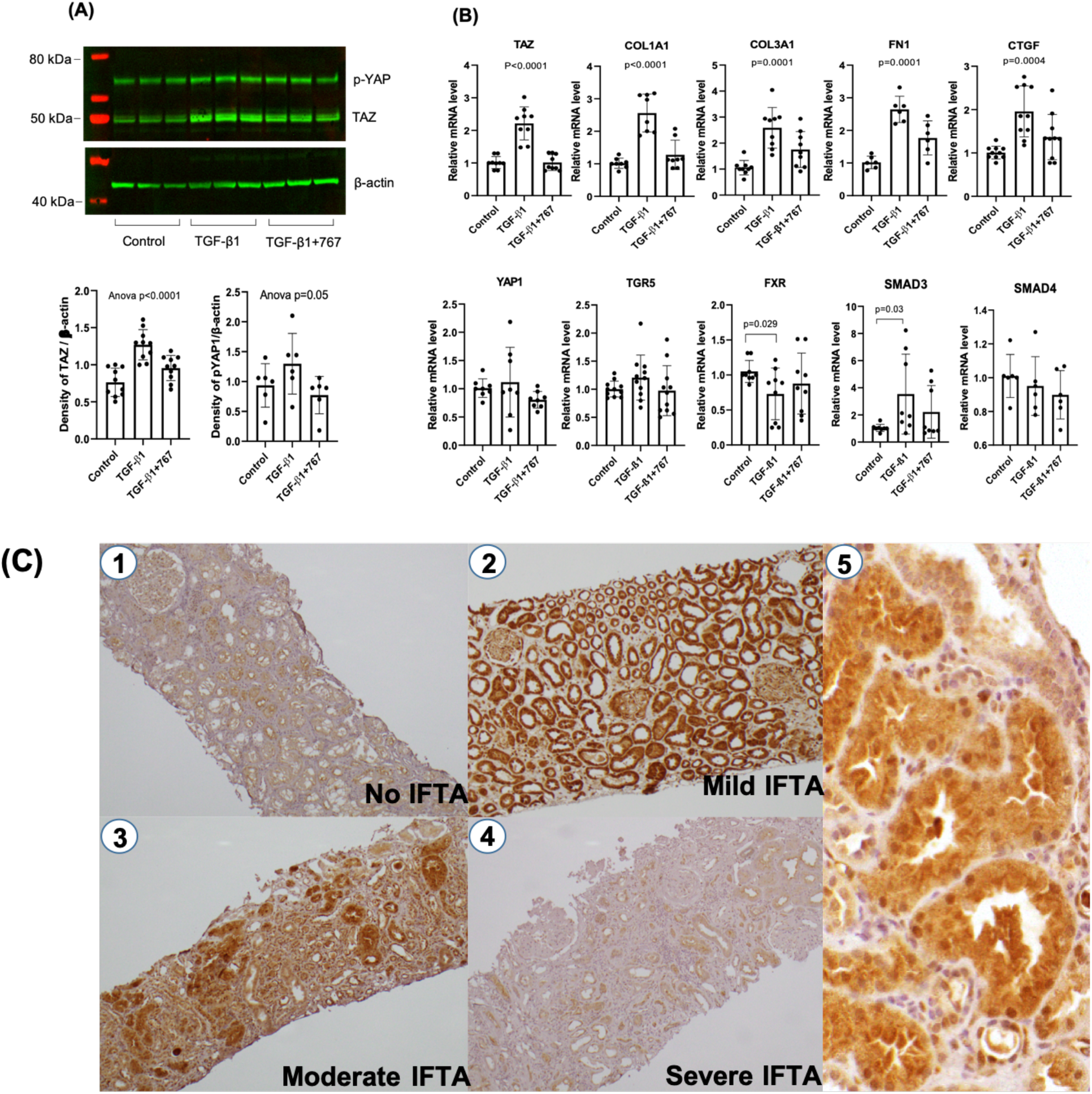
Fibrotic genes and protein expression in kidney organoid, TAZ staining in mild/moderate acute tubular injury. (A) Western blotting showed increase of TAZ protein in organoids exposed to TGF-β1, with band intensity of p-YAP and TAZ proteins normalized to β-actin shown in the dot-plots. The addition of INT-767 inhibited the increase. p-YAP1 protein manifested no significant change. Data are from three independent experiments and are expressed as mean ± SD analyzed by one-way ANOVA. (B) RT-qPCR was used to evaluate mRNA expression of the Hippo pathway transcription factors TAZ and YAP1, as well as FXR, TGR5, SMAD3 and 4, and extracellular matrix proteins. Results were obtained from three independent experiments, normalized to GAPDH expression. Relative mRNA values were calculated by 2^−ΔΔ*CT*^ method and are expressed as mean ± SD. The p value < 0.05 indicates statistical significance by one-way ANOVA. TGF-β1 induced mRNA increases in *TAZ, COL1A1*, collagen 1A3 (*COL1A3*), fibronectin (FN1), connective tissue growth factor (CTGF). These increases were attenuated in the presence of INT-767. TGF- β1 also induced mRNA increase in SMAD3 but decrease in FXR. TGR5, and YAP1, and other trans cription factors of SMAD4, and TEADs (data not shown) remained unchanged. (C) Kidney biopsies obtained from subject with acute tubular injury and variable interstitial fibrotic expansion, immunohistochemically stained for TAZ. Panel 1 shows minimal interstitial fibrosis and tubular atrophy (IFTA), with weak TAZ staining (100X). Panel 2 shows mild IFTA with diffuse intense tubular TAZ staining (100X). Panel 3 shows moderate IFTA with diffuse and intense tubular TAZ staining (100X). Panel 4 shows severe IFTA with weak and focal TAZ staining (100X). Panel 5 shows higher magnification image demonstrating cytoplasmic and nuclear TAZ staining pattern (400X).

TGF-β1 treatment increased mRNA expression of TAZ and the fibrotic genes *COL1A1* (collagen 1α1), *COL3A1* (collagen 3α1), *FN1* (fibronectin), and *CTGF* (connective tissue growth factor). Addition of INT-767 together with TGF-β1 moderated these increases. The extent by which INT-767 reduced the increases in mRNA was as follows: for TAZ, 54%, p<0.0001; for COL1A1, 50%, P<0.0001; for COL3A1, 32% p=0.0001; for FN, 33%, p=0.0001 and for CTGF, 30%, p=0.0004, all tested by ANOVA (**Figure 5B**).

TGF- β1 also induced increase of SMAD3 mRNA (P=0.03) but decrease in mRNA encoding FXR by 30% (p=0.03), (**Figure 5B**). RNA expression of other transcription factors, *SMAD4* and *TEAD* family members, was not statistically altered by treatment.

The time course of downstream effects of TGF-β1 exposure, reflecting couple signaling pathways. An increase in p-SMAD3 occurred within the first 2 hours following TGF-β1 exposure and was sustained for 72 hours (**Supplemental figure 1**). TAZ expression increased after 8 hours of TGF-β1 exposure, TEAD protein levels changed much later; starting to rise 24 hours after TGF-β1 exposure. These effects, once induced, were sustained for up to 72 hours in the presence of TGF-β1. In addition to INT-767 significantly inhibiting the TGF- β1-induced increase of p-SMAD3 protein (**Figure 4B, Supplemental Figure 1**), INT-767 also inhibited the induction of TAZ (**Figure 5, Supplemental Figure 1**).

### TAZ expression is increased in early but not late fibrosis in human kidney

We next sought to establish whether changes of TAZ expression in renal organoids was relevant to human kidney fibrogenesis. To this end we immunostained 44 kidney biopsies with acute tubular injury, in the following fibrosis categories: 7 biopsies with no/minimal fibrosis, 9 with mild fibrosis, 17 with moderate fibrosis and 11 with severe fibrosis. We observed variable nuclear TAZ staining **(Figure 5C)**. In cases with severe acute tubular injury (defined by the presence of cytoplasmic apical blebbing and tubular dilatation and no or minimal interstitial fibrosis or tubular atrophy (<5%) limited TAZ staining was noted (**Figure 5C, panel 1**). In biopsies with acute diffuse tubular injury with diffuse flattening, dilatation of tubules and interstitial edema with mild interstitial fibrosis and tubular atrophy, tubular epithelial nuclear TAZ staining was intense (**Panels 2 and 5**) and similar staining was observed in non-atrophic tubules in biopsies showing more advanced scarring (**Panel 3**). In patients with acute tubular injury and severe tubulointerstitial scarring, only focal and weak TAZ staining was appreciated (**Panel 4**). The cases with the strongest TAZ staining were as follows: 1 out of 7 biopsies in the no/ minimal fibrosis category, 10 out of 26 biopsies in mild/ moderate fibrosis and 2 out of 11 biopsies manifesting severe fibrosis. The proportions of cases manifesting strong TAZ staining cases were 14% in the no/minimal fibrosis group, 38% in the mild/moderate fibrosis group and 18% in the severe fibrosis group (**Figure 5C, Table 1**). Thus, TAZ staining is particularly increased in injured tubules early in the progression of fibrogenesis.

### TAZ interacts with TEADs and p-SMAD3

Tafazzin (encoded by *TAZ)* is a co-transcriptional activator and thus must act in concert with one or more other factors to activate gene transcription (6, 19). TAZ binds the DNA binding protein TEA-domain family member (TEAD) and also acts to shuttle complexes containing SMAD 2/3 or SMAD4 to the nucleus (20). Therefore, we tested whether there is cross-talk or interaction among these three proteins. Immunohistochemical analysis demonstrated that TAZ and TEAD co-localized in renal tubular nuclei within the organoids (**Figure 6A**). TGF-β1 reduced the co-localization, while the FXR/TGR5 dual agonist INT-767 increased co-localization (**Figure 6B)**. Further, we evaluated whether TAZ binds p-SMAD3 and TEADs. Indeed, in HK2 cells, co-immunoprecipitation showed that TAZ interacted with p-SMAD3 and TEADs (**Figure 6C**), indicating that p-SMAD3, TAZ and TEADs may co-regulate fibrotic gene transcription in the kidney.

**Figure 6.**
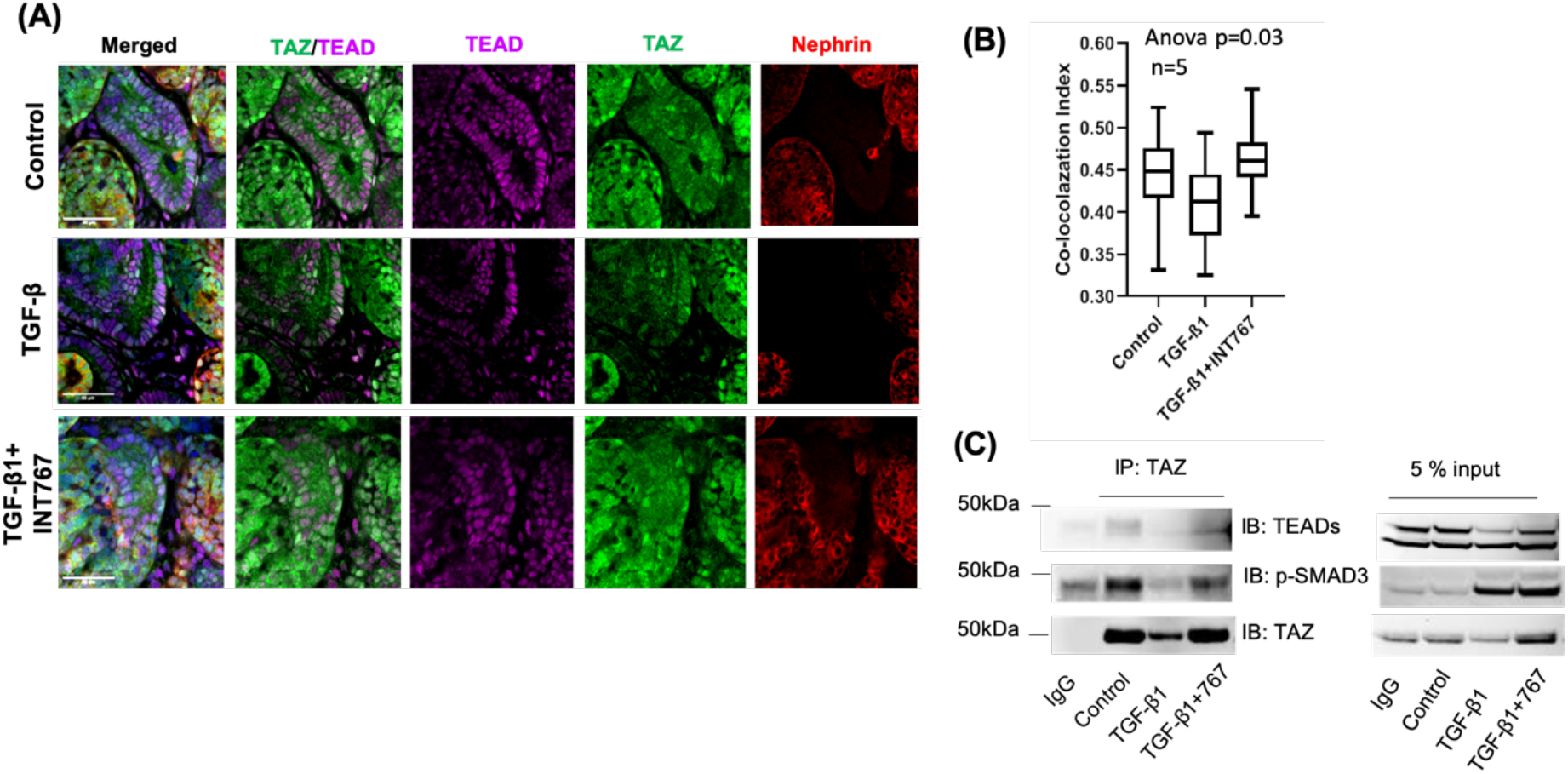
TAZ/TEAD/pSMAD3 co-localization in renal tubules. (A) TAZ and TEAD co-localization. When immunostaining whole organoid with anti-TAZ and TEAD antibodies, the organoids showed TAZ and TEAD co-localization in nuclei, most prominently in renal tubules; green, TAZ; purple, TEAD; red, nephrin. TAZ and TEAD co-localization is shown in white. (B) Quantification of TAZ/TEAD co-localization. The coefficient of two-color co-localization was expressed as co-localization index the significance among the groups is determined by one-way ANOVA. (C) TAZ interacted with pSMAD3 and TEADs. Immunoprecipitation (IP) of HK2 cell lysate showed that TAZ pulled down p-SMAD3 and TEADs, indicating that these three proteins might form a transcription complex. TAZ only pulled down larger TEAD molecule not smaller TEAD molecule (left side panel).

### Intranuclear *TAZ interacts with TEAD1/TEAD4 isoform1 or TEAD1 in renal tubular epithelium*

To determine which TEAD protein interacts with TAZ to promote renal fibrogenesis, we searched GenBank (21) and TISSUES (Tissue expression database, Tissue 2.0 http://tissues.jensenlab.org/), with a focus on two TEAD proteins, TEAD1 and TEAD4 that are highly expressed in kidney. There are three major TEAD4 isoforms. TEAD4 isoform 1 is predicted to have a similar molecular weight as TEAD1 (~50 kDa), and TEAD4 isoforms 2 and 3 have expected molecular weights of ~45 and 35 kDa, respectively. Using a TEAD antibody (Abcam, Cambridge, UK, ab197589) with detects both TEAD1 and TEAD4 proteins, we observed two bands (~50 and ~45 kDa) in renal tubule epithelial cells (HK2) lysate (**Figure 7A, Figure 6C Right**).

**Figure 7.**
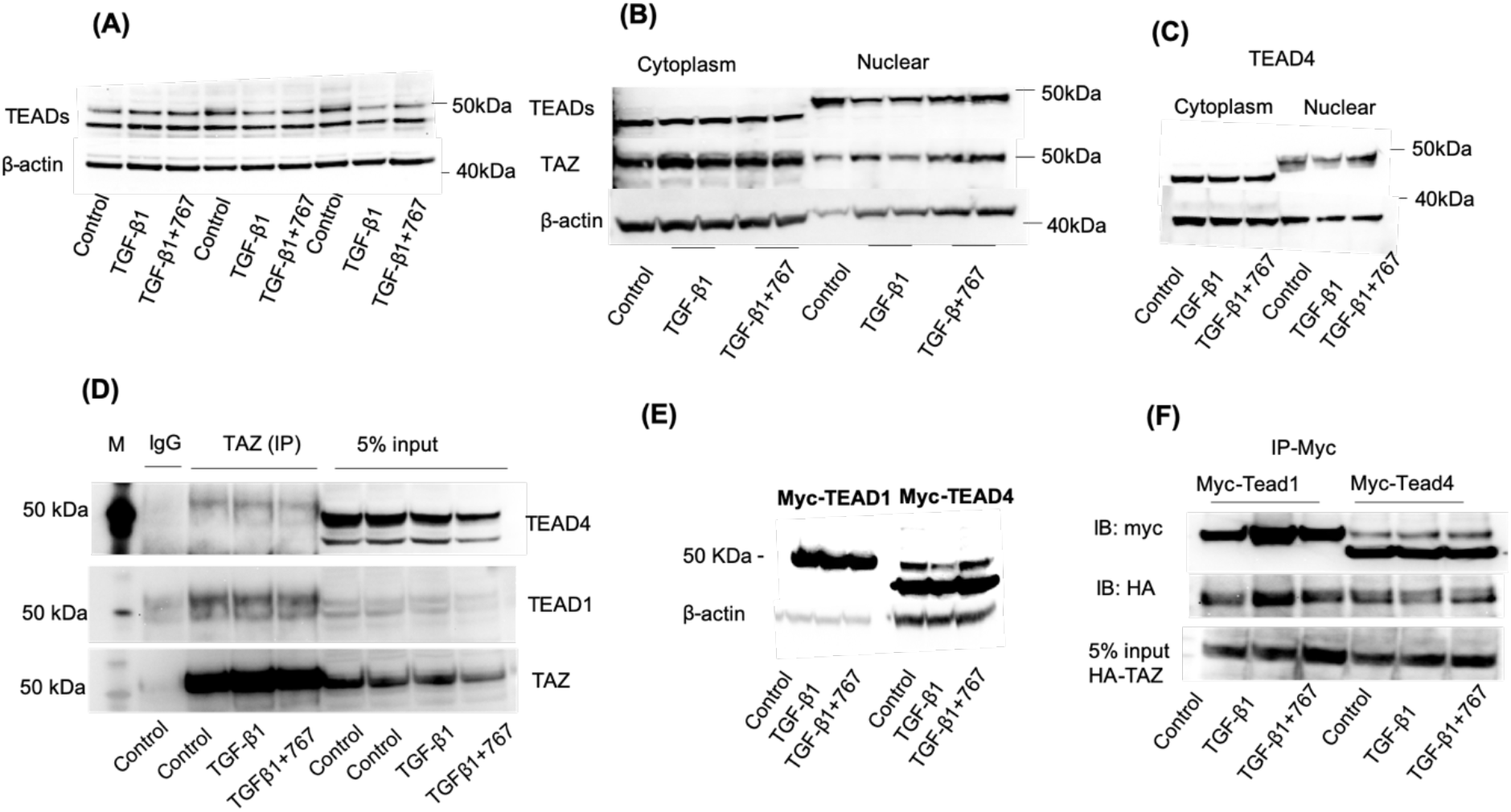
TGF-β changed TEAD nuclear distribution in HK2 cells. TAZ interacted with TEAD4 nuclear isoform (isoform 1) and TEAD1. HK2 cells were used to study the role of TGF-β1 in regulating TEAD cellular distribution. (A) TEAD immunoblot shows two bands TEAD1 and TEAD4 isoform 1 (~ 50 kDa) and TEAD4 isoform 2 (~45 kDa) expression. (B) Cytoplasmic and nuclear fractionation demonstrates that the lower molecular weight TEAD (~45 kDa) was located principally in the cytoplasmic fraction (left panel) and the larger TEAD band (composed of TEAD1 and TEAD4) was located primarily in nuclear fraction (right panel). TGF-β1 exposure reduced TEAD1/4 in the nuclear fraction without affecting the cytoplasmic forms. (C) Immunoblot with TEAD4 antibody confirms TEAD4 isoform 1 (~50 kDa) in nuclear fraction and isoform 2 (~45kDa) in cytoplasmic fraction with reduction of TEAD4 isoform 1 by TGF-β1 (D) TAZ interacted with TEAD4 isoform1 and TEAD1, as shown by TAZ pull-down. TAZ bound TEAD1 and TEAD4. HK2 lysates immunoprecipitation showed that TAZ interacted with the TEAD1 and TEAD4 of nuclear form (isoform 1) but did not bind to theTEAD4 cytoplasm form (isoform 2, right panel, low molecular band). (E) Western blotting of over-expressed myc-TEADs and HA-TAZ in 293T cells. There were a single TEAD1 molecular form and two TEAD4 isoforms with different molecular weights. TGF- β1 exposure reduced the amount of the larger TEAD4 protein. (F) TAZ bound myc-tagged TEAD1 and TEAD4. 293-T cells were transfected with tagged vectors, HA-TAZ plus myc-TEAD1 or myc-TEAD4. At 48 h after transfection, cell lysates were used for myc-immunoprecipitation assay to confirm TEAD1/4 interaction with TAZ. The result showed the myc-TEAD1 or myc-TEAD4 pulled down HA-TAZ (upper 2 panels); increased amounts of HA-TEAD1 and decreased amounts of HA-TEAD4 in the TGF-β1-exposed sample (upper 2 panels).

We hypothesized that the different TEAD isoforms might distribute in different intracellular compartments. Using a celI fractionation approach in HK2 cells, we determined that the larger TEAD molecule (~50 kDa) was present in the nucleus, whereas the smaller TEAD molecule (~45 kDa) was present in the cytoplasm (**Figure 7B**). Moreover, using an anti-TEAD4 antibody, we confirmed that TEAD4 isoforms distribute in different cell fractions. The isoform 1 was located in the nucleus and isoform 2 primarily was in cytoplasm of HK2 cells (**Figure 7C**). However, we did not observe isoform 3 in cytoplasm, as previously reported (19). This is the first observation that TEAD4 isoforms were differentially localized in intracellular compartments in renal tubular epithelial cells.

To investigate possible interaction between TAZ and TEAD proteins, we performed immunoprecipitation experiments in renal tubular epithelial cells. In TAZ pull-down experiments, we found that TAZ interacted with both TEAD1 and TEAD4 isoform 1 (**Figure 7D**) in HK2 cell lysate, as TAZ pulled down TEAD1 and TEAD4 isoform1 but not isoform 2 (**Figure 7D**, right). We demonstrated that TEAD4 isoforms 1 and 2 were present in another renal tubule epithelial cell line, HEK 293-T (**Figure 7E**). TAZ and TEAD1/TEAD4 interactions were also confirmed in HEK 293-T cells overexpressing tagged TEAD1, TEAD4 and TAZ cDNAs. A co-immunoprecipitation experiment showed that myc-TEAD1 or TEAD4 pulled down HA-TAZ protein (**Figure 7F**).

### TEAD4 is a repressor of the collagen1 α1 (COL1A1) promoter and repression is attenuated by interaction with TAZ

In the Eukaryotic Promoter Database (https://epd.epfl.ch//index.php), we found predicted SMAD and TEAD co-binding sites within the *COL1A1* promoter (**Figure 8A)**. We generated a 1 kb *COL1A1* promoter luciferase reporter construct, together with a similar construct but carrying mutations in the potential TEAD binding sequence (**Figure 8A**). We transfected human embryonic renal tubular cells (293-T) with promoterless’ control vector or wild type or the TEAD binding sequence mutant *COL1A1* promoter constructs, together with myc-*TEAD1* or *TEAD4* and HA-*TAZ* expression vectors, and evaluated reported gene activity by measuring luciferase signal. The native-type *COL1A1* promoter had ~3,000-fold higher activity compared to the promoterless’ control vector. Overexpression of TEAD4 repressed the activity of wild type *COL1A1* promoter but did not alter the activity of the mutant promoter (**Figure 8B)**, suggesting this TEAD binding sequence may act as a TEAD4 binding site to repress gene transcription.

**Figure 8.**
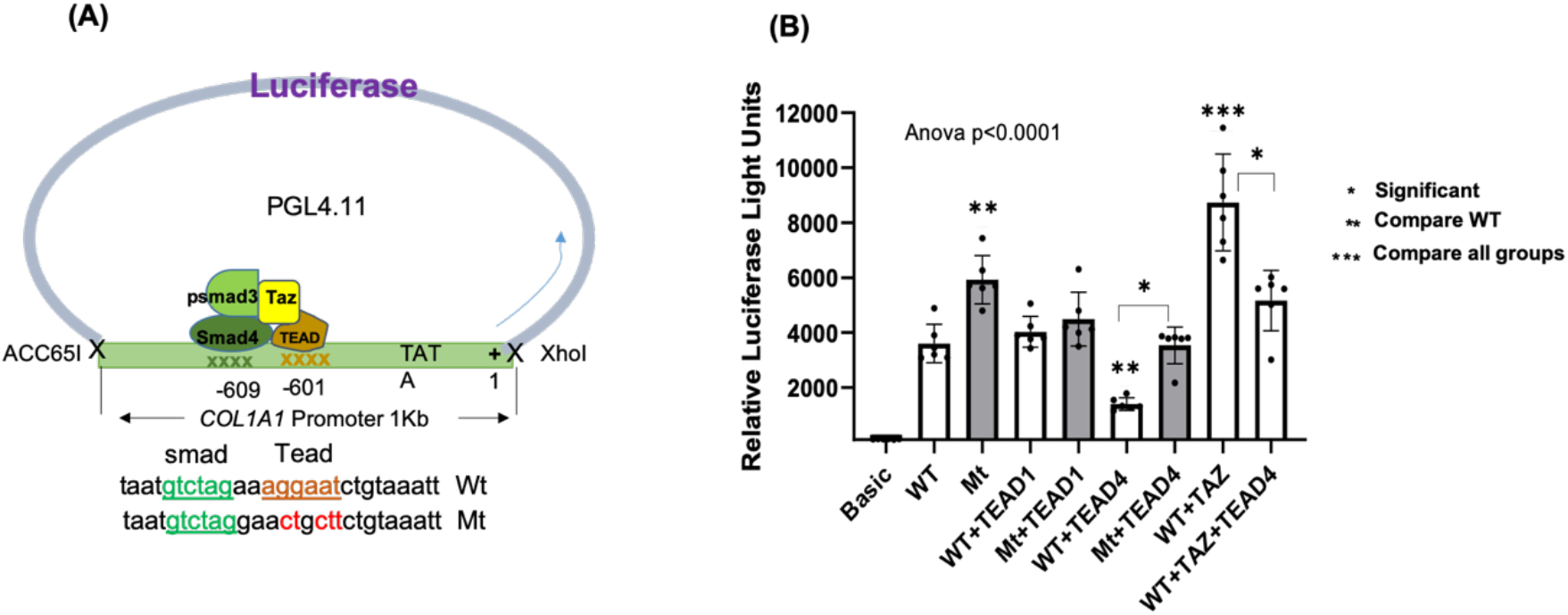
TEAD4 as a transcriptional repressor of the *COL1A1* promoter; TAZ might attenuate TEAD4 repression function. To determine how TEADs and TAZ co-regulate fibrotic gene transcription, we constructed a human *COL1A1* promoter reporter that included a potential TEAD binding sequence. (A) Schematic of two reporter gene constructs. A one kb *COL1A1* promoter was inserted into PGL4.11 basic vector using ACC65I and *Xho*I restriction enzyme sites. A potential TEAD binding sequence GATTCC (compared to the classic sequence CATTCC) is underlined in brown letters, and a potential SMAD (CAGA box) binding sequence is underlined in green letters. The mutated bases are shown in red letters. We hypothesized that SMAD - TEAD-TAZ compound bound to – 601 to 609 bp of the promoter, which includes potential TEAD and SMAD binding sequences. (B) *COL1A1* Luciferase assay. The WT and mutant type (Mt) promoter constructs and transcription factors (myc-TEAD4, myc-TEAD1 or HA-TAZ) as well as the normalizing gene, β-galactosidase, were transfected into 293-T cells, and after 48 h transfection, cells were lysed for luciferase assay. Relative Luciferase activity (RLU) is shown by normalizing light units of luciferase to β-galactosidase units. Values are expressed as mean ± SD. The result shown comprise three independent experiments, each experiment performed in triplicate. ANOVA followed by Turkey multiple comparison with appropriate adjustment to determine the statistics significance between groups. One or more asterisks indicates p<0.05 for comparisons as shown.

**Figure 9.**
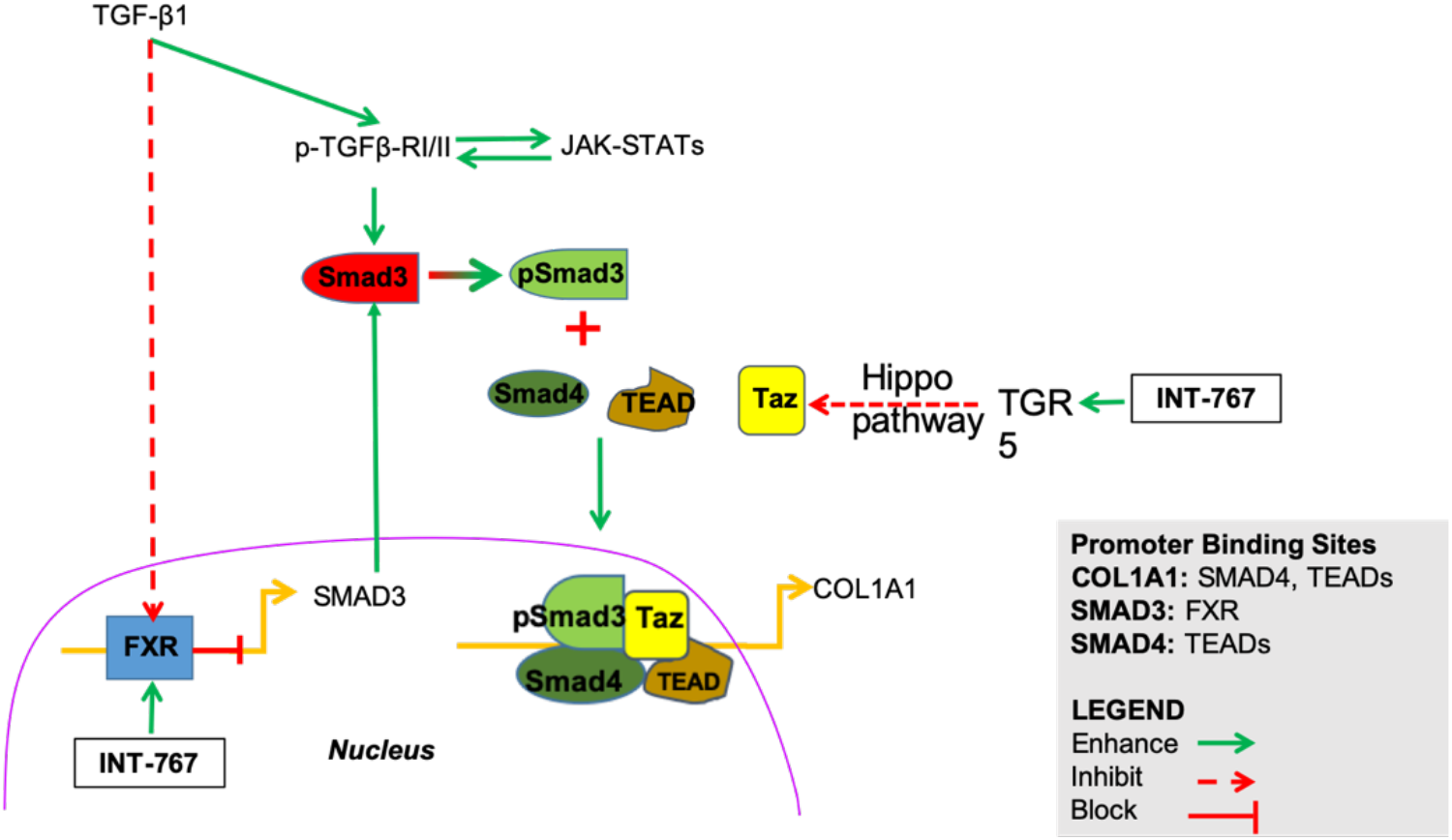
Model of bile acid receptor regulation of the TGF-β1 pro-fibrotic pathway. TGF-β1 activates canonical pro-fibrotic pathways by binding to and signaling through cell-surface receptors, TGFβ-RI/II. These activated (phosphorylated) TGF-β receptor tyrosine kinases phosphorylate SMAD3, with phospho-SMAD3 binding SMAD4. This SMAD hetero-complex translocates to nuclei to activate gene transcription of many genes, some of which are pro-fibrotic. Exposure to TGF-β1results in reduced FXR, a transcriptional repressor of SMAD3. The G-protein coupled bile acid receptor (TGR5) activation inhibits the Hippo pathway. Decreased TAZ phosphorylation and/or de-phosphorylated TAZ may promote shuttling of p-SMAD3 to nuclei, where TAZ binds TEAD transcription factors to initiate target gene transcription. The bile acid receptor agonist INT-767 activates TGR5, suppressing TAZ activation and increases nuclear FXR expression, repressing SMAD3 transcription. Thus, bile acid receptor agonism plays a role downstream of TGF-β1 signaling to block a pro-fibrogenic transcriptional program.

In contrast to the effects of TEAD4, TEAD1 overexpression did not significantly change the TEAD mutant promoter activity, indicating this TEAD binding site in *COL1A1* favors TEAD4 binding over TEAD1. Overexpression of TAZ increased promoter activity compared to the control and co-transfecting TAZ with TEAD4 attenuated the repressive effect of TEAD4 (**Figure 8B**).

## Discussion

The first report of human kidney organoids generated from iPSCs came from Takasato and colleagues in 2015 (22). Recently several groups have used kidney organoids to study inherited kidney diseases, such as congenital nephrotic syndrome due to nephrin mutations (23) and polycystic kidney disease (24, 25); podocalyxin knock out (26); and renal toxicology (27).

There are at least four laboratory protocols describing methods and results of studies using kidney organoids (17, 27–29). Here, we used a modified Takasato protocol. This iPSC culture system is relatively simple, does not require feeder cells, and upon differentiation and aggregation of the cell pellet, the cells form a large kidney organoid. In this study, we used these iPSC-derived organoids to model kidney fibrosis. The mature organoids included diverse cell types that are present in the nephron and associated vascular structures. TGF-β1 exposure triggered a profibrogenic program with extracellular matrix deposition. This adds a new profibrotic trigger, beyond IL-1β (30), to the toolkit of experimenters who wish to model renal fibrogenesis in an in organotypic in vitro system. Further, we used this model to confirm that the FXR /TGR5 agonist, INT-767, attenuates a profibrotic gene expression program and preserves normal nephron structure. These data validate the use of organoids to study kidney fibrosis with numerous advantages, including the ability to study the effect of human genetic background, the potential for scalability and thus a role in screening of chemical and biologic pharmaceuticals.

TGF-β pathways have extensive crosstalk with the Hippo pathway (7) and both contribute to kidney fibrosis (8, 9). YAP/TAZ are major components of the Hippo pathway and serve as co-regulators to activate or repress target genes (6). YAP and TAZ are paralogs, with similar domain structures, and are both similarly regulated by Hippo kinases (6). Interestingly, YAP and TAZ appear to have discrete functions, as suggested by different protein-protein interaction patterns (6, 31). The two factors regulate expression of overlapping similar but distinct sets of target genes. For example, Yap knock-out mice are embryonic lethal with severe developmental defects, whereas Taz knockout mice are viable, with defects in the kidney and lung (32, 33). Angora et al. (34) studied three kidney injury mice models, including CKD, unilateral ureteral obstruction (UUO), and aristolochic acid nephropathy (AAN) and found that TAZ protein levels were markedly increased in all of these models.

We found that expression of TAZ protein but not YAP protein changed significantly in TGF-β1 induced fibrosis in renal organoids, and that TAZ increased in early stages of fibrosis human kidneys. Taken together, these data suggest that TAZ may be an important mediator of dysregulated kidney repair. In response to TGF-β1 stimulation, TAZ shuttled p-SMAD3 to the nucleus (20) with p-SMAD3 bound to various transcription factors to regulate target genes. Co-immunoprecipitation demonstrated that TAZ pulled down TEAD and p-SMAD3, suggesting that the TAZ interaction with p-SMAD3 and TEAD could potentially activate fibrotic genes in renal organoids. In the setting of acute tubular injury in human, we confirmed increased TAZ expression in early stages of kidney fibrosis, but not in advanced fibrosis.

Like its paralog YAP, TAZ binds nuclear transcription factors in order to regulate target genes. TAZ has a TEA-binding domain that allows it to bind TEAD transcription factors. TAZ binding TEADs may enhance or repress the gene transcription, depending on cell type, the particular signaling pathway, and the particular TEAD variant (7, 19). In the present study, TEAD4 acted as a transcriptional repressor of the *COL1A1* promoter and following overexpression, TAZ interacted with the nuclear isoform of TEAD4 and attenuated TEAD4 repression function. This new finding provides a potential target for future investigation into the role of particular TEAD proteins in regulating kidney fibrosis.

In recent years, bile acid receptor agonists have been shown to be effective in treating fibrosis in mouse models of nonalcoholic fatty liver disease and pulmonary disease (35, 36), as well as nephropathy in diabetes, obesity and aging in mice (14, 15). An FXR/TGR5 dual agonist (INT-767) has been shown to prevent progression of nephropathy and reversed kidney injury (14, 15). The therapeutic benefit of INT-767 may come from its multiple actions, including an increase in mitochondrial biogenesis, diminished TGF-β signaling, reduced inflammation, and improved lipid metabolism (35). However, the biological mechanism of bile acid receptor inhibition of TFG- β1 signaling remains elusive. This study shows that INT-767 reduced p-SMAD3 entry into nuclei and TAZ translocation. These data provide mechanisms for the antifibrotic effects of bile acid receptors and suggest new therapeutic strategies.

In summary, this report adds to the literature showing that organoids to be a useful to model study kidney fibrosis and we used this approach to dissect TGF-β1 signaling, including transcription factors SMAD3, TAZ and TEAD4. INT-767, a dual bile acid receptor FXR and GPCR TGR5 agonist, increased FXR expression and TAZ activity, while decreasing p-SMAD3 nuclear translocation, and GPCR-Hippo signaling. The net effect of INT-767 was to prevent fibrosis and to preserve kidney organoid architecture. Future studies are needed to clarify crosstalk between FXR and p-SMAD3, and decipher the pathways linking TGR5 activation and Hippo signaling.

## Methods

### Cell culture and organoid generation

iPSCs were maintained in culture in vitronectin (ThermoFisher, Waltham, MA) coated six-well plates with complete essential 8 medium (E8) (ThermoFisher). The clones were passaged every 4 days at 80% confluency using an EDTA solution-Versene (ThermoFisher). Cells used to generate organoids were between 20-30 passages. Briefly, organoid development occurred in three stages (**Figure 1A)**.

First, cells were plated for differentiation. 80% confluence cell clones were digested with TrypLE Select (Gibco, Gaithersburg, MD), neutralized with E8 medium and centrifuged at 400xg for 3 min. The pelleted cells were re-suspended in E8 medium and ~375,000 cells were seeded into a 25 cm^2^ vitronectin-coated flasks.

Second, intermediate mesoderm cells were generated. The next day, cells reached 40-50% confluence and were placed in E8 medium with APEL2 medium (Stem Cell Technologies, Vancouver, Canada), supplemented with 8 μM CHIR99021, a GSK3 inhibitor (17) to induce WNT signaling for 4-5 day after which cells were cultured in APEL2 medium supplemented with 200ng/mL FGF9 and 1 μg/mL heparin continually for 7 days.

In the final experimental step, aggregated cell pellets were formed. Cells were detached with 0.05% trypsin EDTA, centrifuged and re-suspended. For each organoid, 7⨯10^5^ cells were placed into a 1.5 mL micro-centrifuge tube and centrifuged to form cell pellets. Pellets were transferred to six-well Transwell plates (Corning, Corning, NY) and filled with 1.2 mL APEL2 plus FGF9 and heparin for each well; changed medium every 2 days and withdrew FGF9/ heparin at day 5; the organoids were continually cultured in APEL2 medium to maturation in 25 days.

HK2 cells (ATCC, Manassas, VA) were cultured with a keratinocyte serum free medium (K-SFM, ThermoFisher, Catalog # 17005-042). 293-T cells were cultured with DMEM medium plus 10% FBS. All cell culture was in antibiotic-free conditions.

### Kidney organoid fibrosis model

TGF-β1 was purchased from R & D Systems (Minneapolis, MN; catalog # 240-B-002). INT-767 (MedChemExperess, NJ; Catalog # 1000403-03-1), was dissolved in DMSO as 40 μM stock and stored at −80°C. To study organoid fibrosis, TGF-β1, INT-767, or TGF- β1 plus INT-767 was added to culture media in a final concentration of 5 ng/mL (TGF- β1) and 20 μM (INT-767), respectively. Additionally, control and TGF-β1-treated organoids also contained 0.025% DMSO. After 48-72 h in culture, organoids were fixed with 4% cold paraformaldehyde, washed with Dulbecco’s PBS (DPBS), suspended in DPBS, and stored at 4 °C for fluorescent staining.

### Immunofluorescent staining of whole organoids and frozen sections

Mature organoids on Transwell inserts were fixed with 4% cold paraformaldehyde in 4°C for 2 h and washed three times with cold 1X DPBS. Each organoid was cut out from filter membrane and blocked with blocking buffer (10% donkey serum and 0.3% TritonX-100 in DPBS) at room temperature for 2.5 hours. Diluted primary antibody with blocking buffer was added to the organoid and incubated at 4°C overnight. The organoid was washed with DPBS, 3 times for 30 minutes at room temperature and once overnight at 4°C. The next day, secondary fluorescent antibody in antibody dilution buffer (2% BSA, 0.2% Triton X-100 in DPBS) was added to the organoid and incubated at room temperature for 1.5 hours; washed with DPBS 4 times, then 4’, 6-diamidino-2-phenylindole dihydrochloride (DAPI) was added for nuclear staining, mounted with a ProLong-Gold Antifade Mount (ThermoFisher catalog # p36930). Antibodies used are listed in **Table 2**.

**Table 2a.**
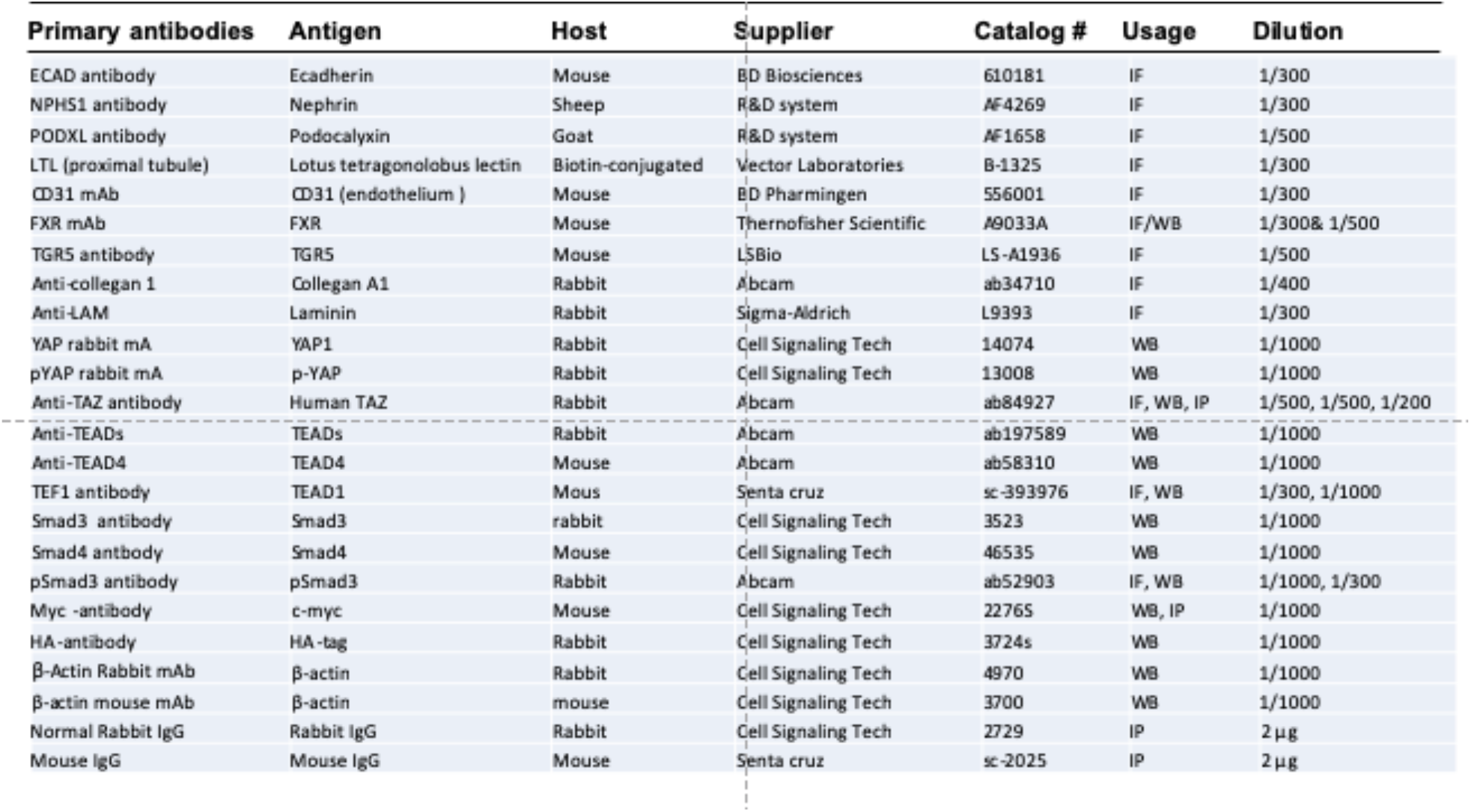
Primary Antibodies.

Paraformaldehyde-fixed organoids were stabilized with 50% sucrose in 4°C overnight and embedded in OCT compound and frozen at −80 °C. Frozen organoids were sectioned at 10 μm for immunofluorescent staining. Fluorescent images were obtained using a Zeiss LSM 700 confocal microscope.

*Cell fractionation*5 ⨯ 10^6^ HK2 cells were seeded into 10 mm plates with K-SM medium. The following day, cells were 70% confluent. DMSO, TGF-β1 + DMSO, or TGF-β1 + INT-767 (20 mM dissolved in DMSO) were added to culture medium in the final concentration of 0.05%, 4ng/mL + 0.05%, 4 ng/mL+ 10 μM, respectively. After 48 hours in culture, cells were used for fraction study. Cell fraction preparation was as previously described (37). Briefly, Cells were washed once with cold buffer A, composed of 10 mM Na-HEPES, pH 7.9, 1.5 mM MgCI_2_, 10 mM KCI, add 1mM DTT and 1 mM phenylmethylsulphonyl fluoride. Cold buffer A, 2 mL, including 0.1% Nonidet P40 and 1% proteasome cocktail (catalog # P8340, Sigma-Aldrich, St Louis, MO) was added to the plates, and cells were scraped into 10 mL tubes and incubated on ice for 15 minutes. The cell suspension was centrifuged at 850xg for 5 minutes at 4°C, and the supernatant as cytoplasmic fraction was collected and concentrated with a Centricon column (10-kDa cutoff, Millipore-Sigma-Aldrich). The pellet as nuclear fraction was lysed with a lysis buffer including 0.1 N NaOH and 0.1 % SDS.

### Co-Immunoprecipitation and immunoblotting

TGF-β1 or INT-767 treated HK2 or 293-T cells were harvested in RIPA buffer (Sigma # R0278); 200 µg cell protein in 400 μL volume was added to 2 µg primary antibody and gently rotated at 4°C overnight, and 25 μL of protein-A/G agarose beads (#sc-2003, Santa Cruz Biotechnology, Dallas, TX) was added to each reaction and rotated for two h at 4^0^C. After four cycles of washing and centrifugation, the pellet was re-suspended, denatured, and run on 4-12 % or 10% NuPAGE gel (Thermofisher Scientific) for immunoblotting with respond antibodies. To avoid IgG bands masking proteins of interest in IP samples, a TidyBlot conjugated Western blot detection reagent (BioRad, Hercules, CA) was used which bound to native non-denatured antibodies and not to any denatured IgG present in the immunoprecipitation samples. Immunoblotting procedure as previous described (37). The membranes were stripped with a strip buffer (Thermofisher Scientific, Waltham, MD) and re-blotted as needed. The blotting images were obtained using an iBlot 2 system (Thermo Fisher Scientific). Antibodies for immunoblotting are summarized in **Table 2a and b**.

**Table 2b.** Secondary Antibodies.

| Secondary antibodies | Supplier | Catalog # | Usage | Dilution |
| --- | --- | --- | --- | --- |
| Donkey anti rabbit IgG (H+L), Alexa Fluor 488 | Thermofisher Scientific | A32790 | IF | (1:500) |
| Donkey anti-Mouse IgG (H+L), Alexa Fluor 488 | Thermofisher Scientific | A21202 | IF | (1:500) |
| Donkey anti-Goat IgG (H+L) , Alexa Fluor 488 | Thermofisher Scientific | A11055 | IF | (1:500) |
| Donkey anti-Rabbit IgG (H+L), Alexa Fluor 568 | Thermofisher Scientific | A10042 | IF | (1:500) |
| Donkey anti-Mouse IgG (H+L), Alexa Fluor 568 | Thermofisher Scientific | A10037 | IF | (1:500) |
| Donkey anti-Sheep IgG (H+L), Alexa Fluor 568 | Thermofisher Scientific | A21099 | IF | (1:500) |
| Donkey anti-Rabbit IgG (H+L); Alexa Fluor 647 | Thermofisher Scientific | A31573 | IF | (1:500) |
| Donkey anti-Mouse IgG (H+L), Alexa Fluor 647 | Thermofisher Scientific | A32787 | IF | (1:500) |
| Chicken anti-Rabbit IgG (H+L), Alexa Fluor 647 | Thermofisher Scientific | A21443 | IF | (1:500) |
| Streptavidin, Alexa Fluor™ 647 Conjugate | Thermofisher Scientific | S32357 | IF | (1:500) |
| Streptavidin, Alexa Fluor™ 405 conjugate | Thermofisher Scientific | S32351 | IF | (1:500) |
| Streptavidin, Alexa Fluor 488 conjugated | Thermofisher Scientific | S32354 | IF | (1:500) |
| Anti-Rabbit IgG, HRP-linked antibody | Cell Signaling Technology | 7074 | WB | (1:5000) |
| Anti-mouse IgG, HRP-linked antibody | Cell Signaling Technology | 7076 | WB | (1:10000) |

### Gene expression assay

Organoid total RNAs were isolated using Trizol reagent (ThermoFisher), following manufacture’s instruction. Isolated RNA was reversed transcript to first strand cDNA using a SuperScript first strand cDNA synthesis kit (ThermoFisher, Catalog # 11904018). The cDNAs were used for quantitative polymerase chain reaction (qPCR). Primers for qPCR were designed with Primer-BLAST (35). The primers for fibrotic gene qPCR are listed in **Supplemental Table 2**. qPCR was performed in a Quantstudio 6 thermocycler (ThermoFisher) and using the qPCR reagents (Agilent Technologies, Cat# 600882). GAPDH served as normalizing gene and the relative mRNA levels of target genes were calculated by 2^−ΔΔ*CT*^ method, and expressed as mean ± SD.

### Wild type and mutant type COL1A1 promoter constructs and luciferase assay

PGL4.11 basic vectors, β-galactosidase expression vector, and assay reagents were bought from Promega (Madison, WI). The plasmids with cDNA of HA-TAZ, myc-TEAD1 and myc-TEAD4 were obtained from Addgene, Watertown, MA (catalog # 32839, 33109 and 24638 respectively). A 1Kb *COL1A1* promoter fragment (GenBank NC_000017.11) was amplified by PCR with a human genomic DNA (Roche, Catalog# 1169111200). The primers were 5′tc*ggtacc*tcaccaatgatcacaggcctc and 5′ac*ctcgag*aaactcccgtctgctccga including an *Acc*65I and a *Xho*I sites, indicated by the italicized letters.

In order to mutate the potential TEAD binding site, two primer pairs that included TEAD mutant sites were used to amplify two small mutant fragments. The 1 kb mutant TEAD fragment was obtained by combining the mutant fragments with overlap PCR (34). The PGL4.11 basic vector and amplified fragments were digested with *Xho*I and *ACC*65I and inserted by ligation into the PGL4.11 vector upstream of the luciferase cDNA. Positive clones were screened by PCR and confirmed by sequencing. Plasmids were co-transfected into the 293-T cells as needed, using a Lipofectamine 2000 reagent (ThermoFisher Scientific). Cells were harvested and lysed 48 h after transfection and luciferase bioluminescence was measured to asses promoter activity, with relative luciferase light units (RLU) normalized to β-galactosidase activity units.

### Fluorescent staining quantitation

To quantify COL1A1 in slides stained for epifluorescence, the HALO Image analysis software, HALO software version 2.0 (Indica Labs. Albuquerque, NM) was used by an area quantification program - Area quantification FL, which quantifies and reports dye positive area, average intensity for each dye. Extent of COL1A1 staining Quantification was calculated and expressed as percent of total image area. Co-localization measurement was used image J-co-localization analysis program, and the co-localization index indicated two color overlap co-efficiency. Both COL1A1 Quantification and TAZ-TEAD co-localization analyses included five organoids for each group, and for each organoid, 4-5 image sections were analyzed.

### TAZ immunohistochemistry

Deidentified human kidney biopsies from the Johns Hopkins Renal Pathology Archives were used under an IRB-approved protocol. Formalin fixed and paraffin embedded kidney biopsy slides were deparaffinized with xylene for 10 min, three times, then rehydrated with gradient concentrations of ethanol. For antigen retrieval, the slides were transferred into a pressure cooker including a Trilogy solution (Sigma, Catalog No. 920P), boiled under high pressure for 30 min.

For immunohistochemistry, slides were blocked with a peroxidase blocker (Bio SB, Santa Barbara, CA, catalog No. BSB 0054) for 5 min; washed with an inmmuoDNAwasher buffer (Bio SB, catalog No. BSB 0150) once; and blocked with an antibody diluent (Bio SB, Catalog No. BSB 0041) for 20 min. Slides were incubated with 1:100 dilution of rabbit anti-human TAZ antibody (Abcam, catalog No. ab 84927) for 1 hour. After three washes, a high sensitive Mouse/Rabbit PolyDetector Plus DAB HRP Brown Detection System (Bio SB, Catalog No. BSB 0269) was used to develop tissue staining following manufacturer’s instruction. Hematoxylin was used as a counterstain.

### Statistical analysis

All quantitative experiments were repeated at least three times and all quantitative assays were performed in triplicate. Results were expressed as mean ± SD. Student *t* test was used to determine the significance of difference between two groups. One-way ANOVA was used to determine the significance of differences among three or more groups. Turkey test was used for multiple comparisons after ANOVA.

## Author contributions

XPY designed and performed experiments, collected and analyzed data, and wrote the first draft of the manuscript. MD performed immunofluorescence assays and obtained images. PD assisted in using image J program. PF analyzed human biopsy samples and obtained images. KPM analyzed collagen1 (α1) images. XW provided INT-767 reagent and instruction. ST designed partial qPCR primers and performed partial assay; MKH advised on data analysis and edited the manuscript; JBK advised on TGF-β1- induced fibrosis and edited the manuscript. ML advised in experimental design and edited the manuscript. AZR supervised and designed experiments, analyzed data, and wrote and edited manuscript.

## Acknowledgements

This work was supported by the American Society of Nephrology and the Johns Hopkins Startup fund (Avi Rosenberg), NIH RO1 DK116567 and AG049493 (Moshe Levi) and R01HL137811 (Marc Halushka) and NIDDK Intramural Research Program (Jeffrey Kopp). We appreciate Dr. Liming Gou who provided the human stem cell line.

## Figure Legends

**Supplemental Figure 1.**
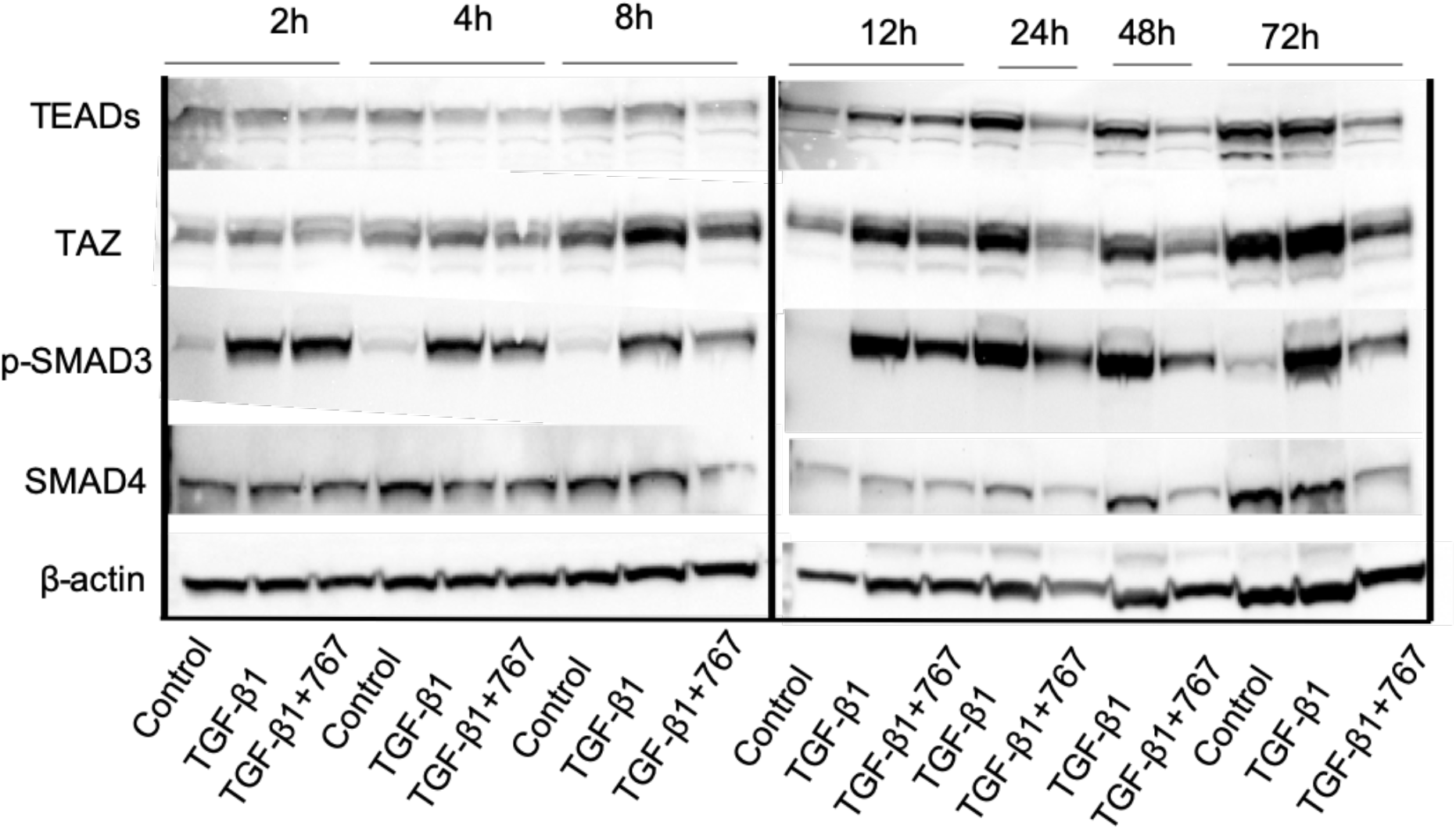
TGF- β1 time course study. TGF-β1 4 ng/mL or the same amount TGF-β1 plus 10 μM INT-767 were added to cultured HK2 cells. TGF-β1 caused a rapid increase in p-SMAD3 within 2h, and this persisted for 72 h; lNT-767 prevented this increase. TAZ increase occurred later, at 8 h and persisted to 72 h, while INT-767 attenuated this increase. TEADs changed much later, starting in 24h after TGF-β1 treatment. In contrast, SMAD4 protein levels remained unchanged.

**Supplement Table 1.**
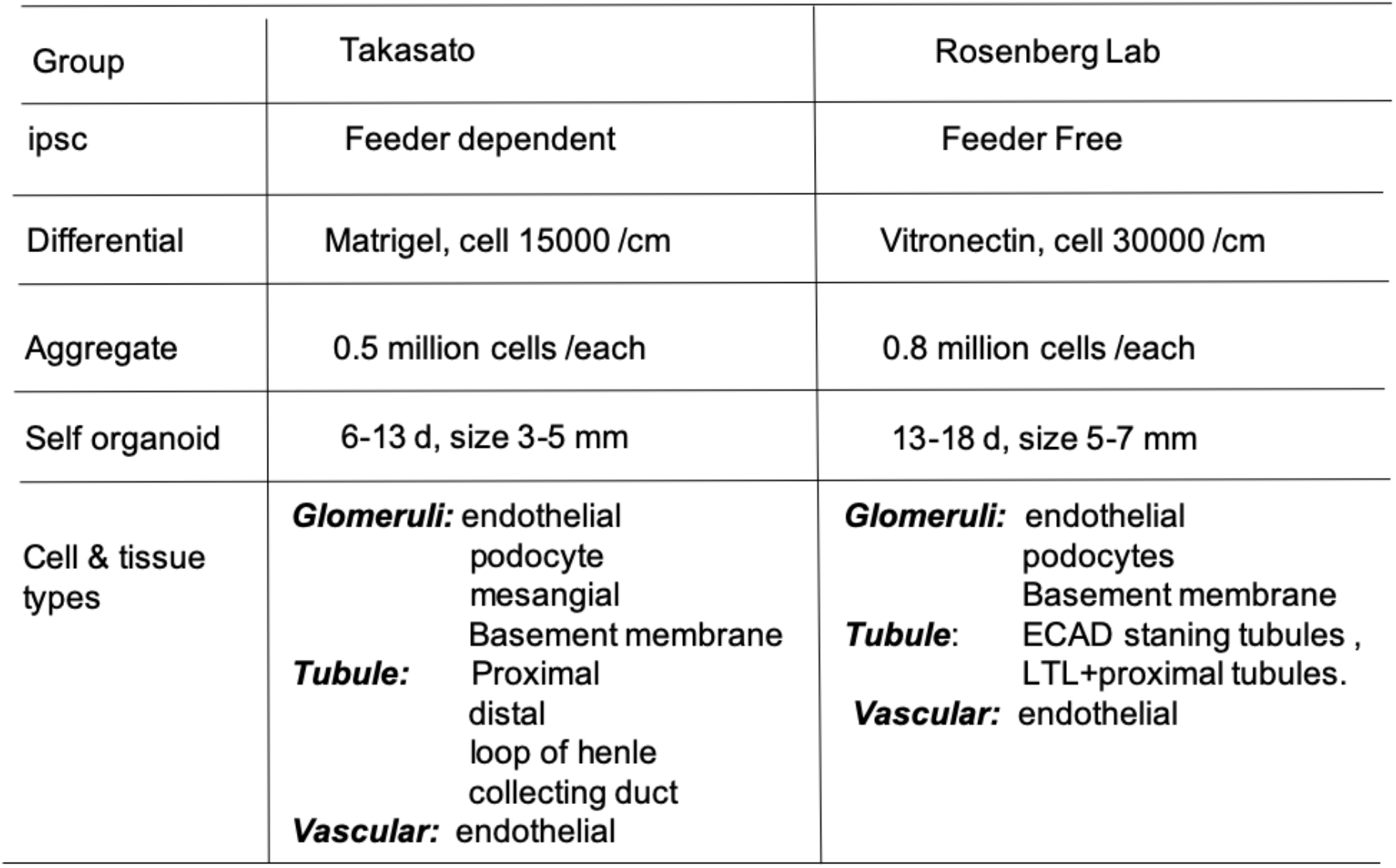

**Supplement Table 2.**
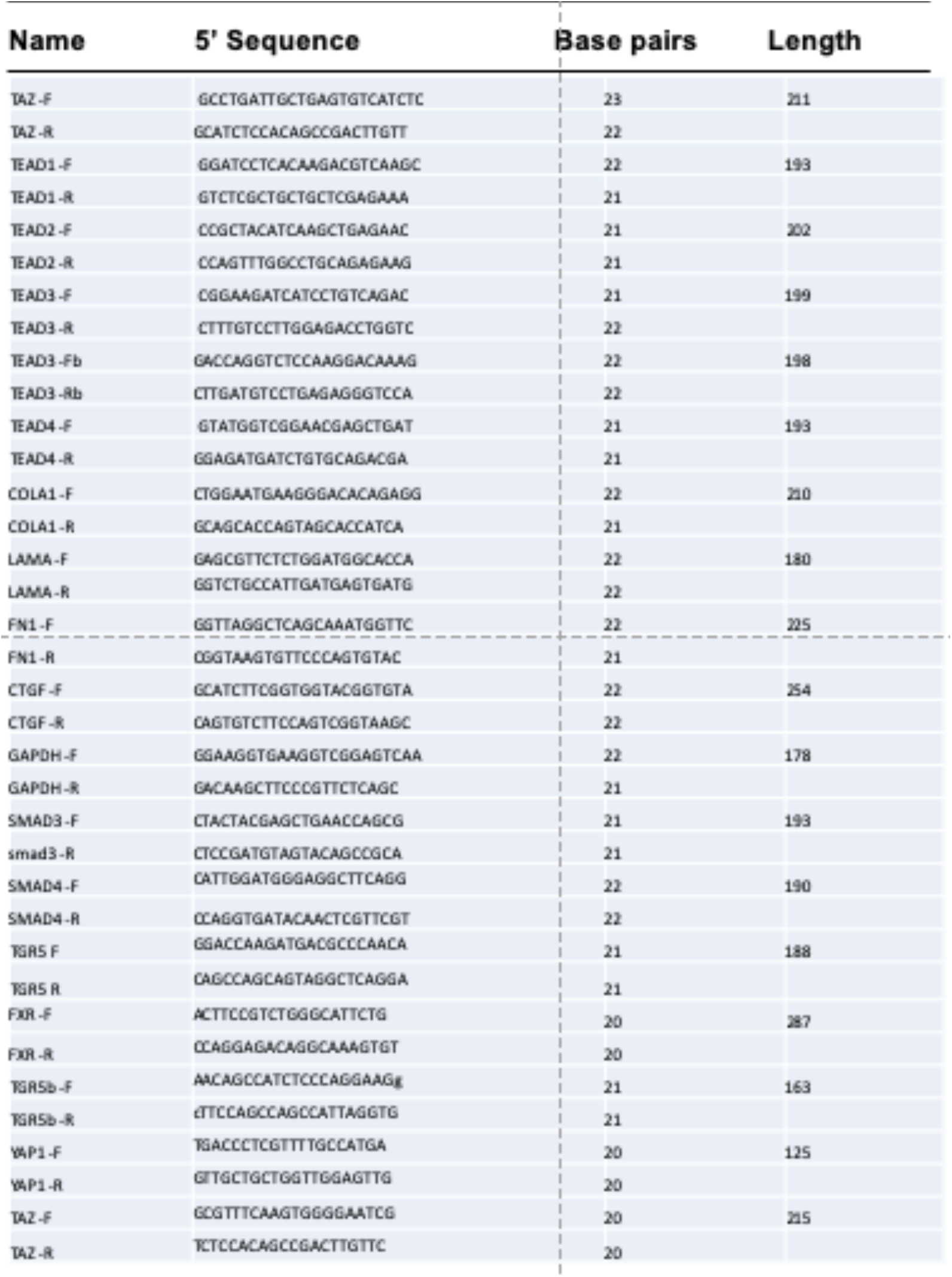

